# Characterization of b-value dependent *T*_2_ relaxation rates for probing neurite microstructure

**DOI:** 10.1101/2022.09.02.506440

**Authors:** Lipeng Ning, Carl-Fredrik Westin, Yogesh Rathi

## Abstract

Brain tissue microstructure is characterized by heterogeneous diffusivity and transversal relaxation rates. Standard diffusion MRI (dMRI) is acquired using a single echo time (TE) and only provides information about heterogeneous diffusivity in the underlying tissue. Combined relaxation diffusion MRI (rdMR) integrates dMRI with multiple TEs to probe the coupling between relaxation rate and diffusivity. This work introduces a method to model rdMRI data signals by characterizing the apparent relaxation rate related to dMRI with different b-values. The proposed approach can extrapolate dMRI signals to ultra-long or ultra-short TEs to increase or reduce signals from intra-neurite water to improve the characterization of neurite microstructure without solving multi-compartment models. The performance of the proposed method was examined using an *in vivo* dataset acquired from a clinical scanner to estimate neurite sizes.

## 1 Introduction

Diffusion magnetic resonance imaging (dMRI) is an imaging technique that probes water diffusion in biological tissue to characterize the underlying microstructure [19]. Different types of microstructural models or signal representation techniques have been developed to characterize the tissue microstructure [2, 26, 8, 18]. But tissue microstructure is not only characterized by hetergeneious diffusion coefficients but also various transversal relaxation rates. The coupling between diffusivity and relaxation rates can lead to echo time (TE) dependent dMRI microstructure [24, 17]. Several methods have shown that integrating dMRI with multiple TEs can better characterize tissue microstructure than dMRI with a single TEs [13, 21, 5, 4].

Current methods for analyzing multi-TE dMRI data can be grouped into two major categories. The first group of methods, such as [24] and [5], use multi-compartment models to characterize signals from different tissue components, i.e., the intra- and extra-neurite signals, with different diffusivity and relaxation rates. These models include more unknown parameters to characterize the *T*_2_ relaxation rates than the corresponding models for single-TE dMRI data. Moreover, they rely on explicitly modeling of both intra- and extra-neurite signals. But solving a multi-compartment models is still generally a challenging problem [7]. The second group of methods focus on characterization of the joint distribution of *T*_2_ relaxation rates and the diffusivity of the underlying tissue components [13, 4, 3] or the moments of the joint distribution [17]. This method typically requires multiple samples of TEs to characterize the under distributions.

In this work, we introduce a new method for modeling multi-TE dMRI data base on b-value dependent relaxation function. Our method first characterizes the apparent *T*_2_ relaxation rate of dMRI signals at different b-values then extrapolate signals to ultra long or ultra short TEs to enhance or reduce signals related to different components. It was shown in [15] that intra-neurite water usually has much longer *T*_2_ than the extra-neurite component. Thus the extrapolated signals at ultra-long TE are dominantly from the intra-neurite water since the underlying extra-neurite component is significantly attenuated at long TE. The extrapolated ultra-long TE signal can be used to neurite microstructure without solving multi-compartment models. The essence of the proposed method is closely related to the work by [25] on probing tissue microstructure using ultra-high b-values. The performance of the proposed method is illustrated using a example on neurite size estimation using a clinical scanner.

## 2 Theory

### 2.1 Signal model

Let *S*(*b, TE*) denote the rdMRI signal at a given b-value and a TE. Existing methods for modeling *S*(*b, TE*) are either based on multi-compartment models or using joint distribution of relaxation and diffusion. In this work, we propose a method to characterize the rdMRI signal around an echo time 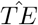 using the following representation

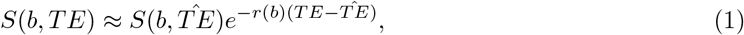

where *r*(*b*) denote the apparent *T*_2_ relaxation rates at different b-values. In practice, values of *r*(*b*) can be estimated using dMRI data acquired at two or more TEs. Then, the above model can extrapolate signals to ultra-long or ultra-short TE values that cannot be directly measured because of restrictions on sequence parameters or limitation on the signal-to-noise ratio.

The model in Eq. (1) provide an approach to push the limit in a different direction, i.e. TE, to estimate neurite microstructure by taking advantage of the different *T*_2_ values between intra and extra-neurite water [15]. Below, we introduce the relation between the b-value dependent relaxation rate *r*(*b*) and multi-compartment models and joint relaxation diffusion distribution

### 2.2 On multi-compartment models

Assume that the rdMRI signal of a voxel is represented by

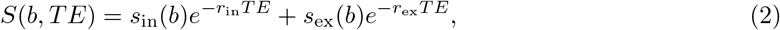

where *s*_in_(*b*), *s*_ex_(*b*) denote the intra-neurite and extra-neurite signals at different b-values and *r*_in_ and *r*_ex_ represent the corresponding *T*_2_ relaxation coefficients. By matching the first order term of *TE* on both sides of Eq. (1), we obtain that

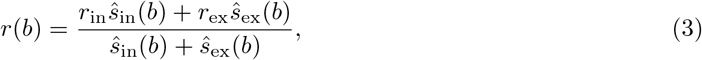

where

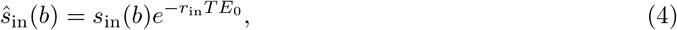

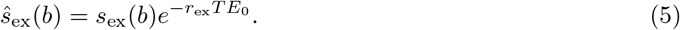

Next, we introduce the following two assumption on the intra- and extra-neurite signals. First, we assume that at *s*_in_(*b*) dominates *s*_ex_(*b*) at large b-values, *s*_in_(*b*) ≫ *s*_ex_(*b*) as *b* → ∞. Second, we assume that the ratio *s*_ex_(*b*)/*s*_in_(*b*) is monotonically decreasing as *b* increases which also indicates more signal contribution intra-neurite water at high b-values.

Based on the first assumption, the following equation holds

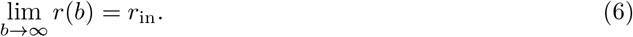

Next, taking the derivative of *r*(*b*) gives rise to

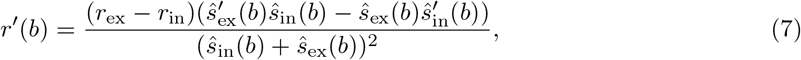

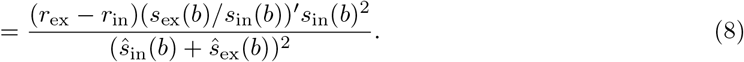

Thus,

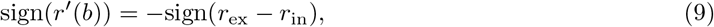

which indicates that *r*(*b*) is a monotonic function. If *r*_ex_ > *r*_in_, then *r*(*b*) is a monotonic decreasing function of *b* and converges to *r*_in_.

### 2.3 On joint relaxation and diffusion distributions

The *r*(*b*) function can also directly reflect the joint moments of the underlying distribution of relaxation and diffusion coefficients. Based on [13, 4, 3, 17], heterogeneous tissue microstructure is model by a joint distribution of relaxation and diffusion coefficients *ρ*(*r, D*) with dMRI signal from each component modeled by *e*^−*bD*−*TEr*^. Then the rdMRI signal is represented by

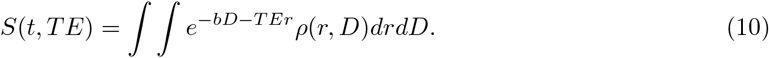

Let (*m, n*)th order central moment of *r* and *D* be denoted by

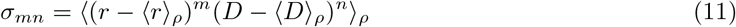

where ⟨·⟩_*ρ*_ denotes the expectation with respect to *ρ*. Then, the cumulant expansion of *S*(*b, TE*) that only contains *σ_mn_* with *m* ≤ 1 is given by [17]

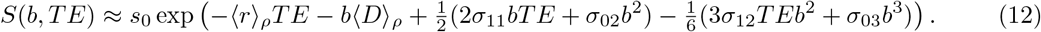

Thus, *S*(*b, TE*) can be represented using 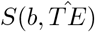 as

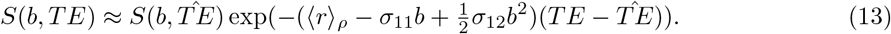

Comparing Eq. (13) with Eq. (1) gives rise to

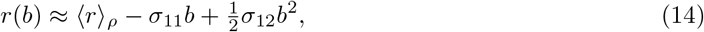

where the higher order terms of *b* is not consider. Thus, −*σ*_11_ and *σ*_12_ are the first and second order derivatives of *r*(*b*) at *b* = 0, respectively. Moreover, Eq. (9) indicates that *σ*_11_ ≥ 0 in most brain tissue.

## 3 Experiment

In this section, we present an application of Eq. (1) on neurite size estimation using rdMRI data acquired on a clinical scanner.

### 3.1 *In vivo* MRI acquisition and processing

A healthy adult was scanned using a Siemens 3T Prisma MRI to acquire a dMRI dataset with to TEs. The imaging parameters are as below: voxel size = 2×2×2 mm^3^, matrix size = 100×100×62, *TE*_1_ = 91*ms*, *TE*_2_ = 121*ms*, TR=4700 ms, *δ* = 15.1 ms, ∆ = 59.2 ms, simultaneous multi-slice = 2, in-plane acceleration =3, 20 co-linear gradient directions at *b* = 1, 3, 3.5, 4, 4.5, 5, 5.5, 6 ms/*μ*m^2^ with additional 4 b0 volumes at each TE. An additional b0 volume at *TE*_1_ = 91*ms* was acquired with the reverse phase-encoding direction, i.e. P≫A. The acquired data was processed using Topup and Eddy method in the FSL toolbox [22] to correct for motions and distortions. Then, the random-matrix-based method [23] was applied to denoise images.

### 3.2 Signal estimation

Based on the exampled data, we first computed the direction averaged signals at each TE to generate the sample data 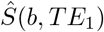 and 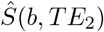. Next, we define the reference 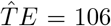 ms as the center point of *TE*_1_ and *TE*_2_ and define

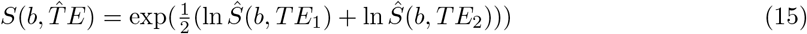

as the geometric mean of the measured data at the two TEs. To estimate the b-value dependent relaxation rate, we first computed

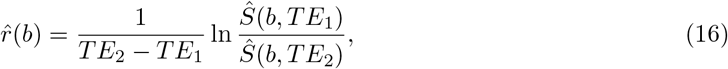

at each b-value. Considering that there are only two TEs which may increase the uncertainlty of the estimated 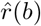, we applied two methods to improve the estimation results by using the property that *r*(*b*) is a monotonic function. For the first method, we fitted 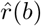 to a model

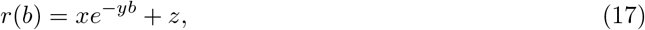

where *x, y, z* ≥ 0. The above model ensures that *r*(*b*) is monotonically decreasing and always takes non-negative values. For the second method, we didn’t use any explicit model for *r*(*b*) and solved a nonlinear least squares (NLS) problem to fit the measurements to exponential functions at each b value with *r*(*b*) being a monotonic decreasing function. Based on the estimated *r*(*b*) values, we applied Eq (1) to predict signals from *TE* = 31 ms to *TE* = 211 ms.

### 3.3 Neurite size estimation

Assume that the diffusion signal from each axon bundle is modeled by a diffusion tensor model with axial diffusivity *d*_∥_ and radial diffusivity *d*_⊥_. Then, the direction-averaged intra-neurite signal is represented by [12]

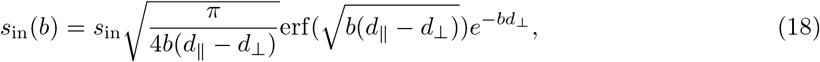

where erf denotes the error function. In [11], *d*_⊥_ was assumed to be zero for intra-neurite signals because of the relatively low b-values used. On the other hand, erf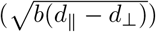 was assumed to be close 1 at ultra high b-values with *bd*_∥_ ≫ 1 in [25]. Here, these simplifications are not imposed in this work because the b-values are much lower than those used in [25] but b = 6 *ms/μm*^2^ is still high enough to probe *d*_⊥_ ≥ 0.015*μm*^2^/*ms* [25].

We applied a NLS fitting method to fit (18) to estimated signals *S*(*b, TE*) at different TEs without explicitly modeling the extra-neurite components. The values of *d*_⊥_ is restricted in the interval 3*μ*m^2^/ms ≥ *d*_⊥_ ≥ 0.015*μ*m^2^/ms and 3*μ*m^2^/ms ≥ *d*_∥_ ≥ *d*_⊥_. Based on the estimated model parameters, we used the following equation from [25] to compute the effective neurite radius

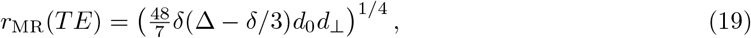

where *δ* and ∆ are the pulse width and diffusion time of the diffusion gradients and *d*_0_ is the diffusivity of the axoplasm which is set *d*_0_ = *d*_∥_ in this experiment. We expect that the value of *r*_MR_(*TE*) reduces as TE increases since there is more intra-neurite signals at long TEs.

## 4 Results

### 4.1 On b-value dependent relaxation

Fig. 1 illustrates the estimated *T*_2_ relaxation maps at different b-values. Overall, the relaxation rate decreases with increasing b-values. Sub-cortical gray matters have the most significant differences between relaxations relates at b=0 and 6 ms/*μ*m^2^. Since extra-neurite signals are approximately vanished at 6 ms/*μ*m^2^, the relaxation rate at 6 ms/*μ*m^2^ is close to *r*_in_. On the other hand, the much higher relaxation rate at *b* = 0 indicating that the relaxation rate of extra-neurite signals are much higher than the intra-neurite signals, i.e. *r*_ex_ < *r*_in_, which is consistent with results from [16].

**Figure 1:**
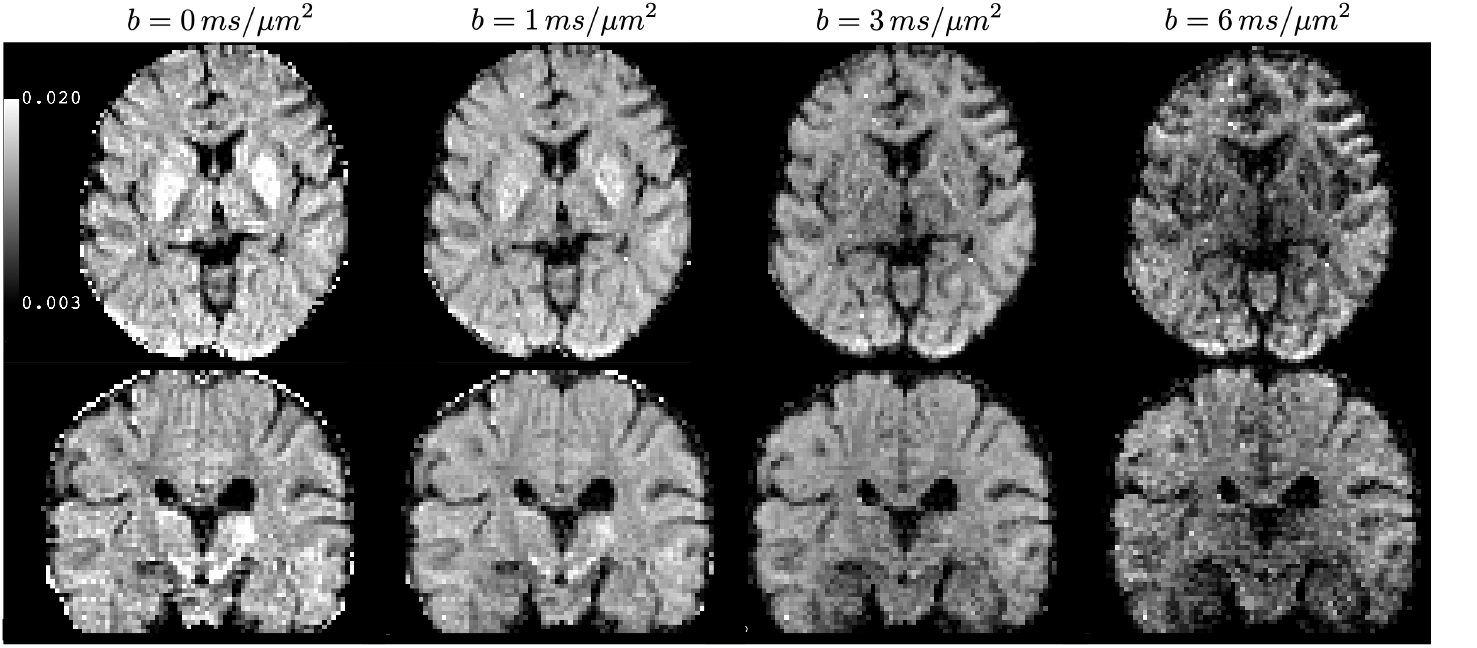
Illustration of the estimated *T*_2_ relaxation coefficients at different b-values with the unit being ms^−1^.

### 4.2 Neurite size estimation

Fig. 2 illustrates the estimated effective neurite size from TE=31ms (left) to TE = 181 ms (right) with *r*(*b*) obtained based on Eq. (17). Overall, the estimated neurite size decreases with increasing TEs because of more intra-neurite signals. At low TEs, the image contrast mainly reflects the difference between gray and white matter. Increased extra-neurite signals at short TEs leads to much larger effective neurite radius. As TE increases, more rich contrast is shown within the white matter and subcortical gray matter. It is interesting to note that the thalamus shows very high radius around 8–19 *μm* at low TEs but with the radii around 1–3 *μm* at long TEs reflecting that the underlying tissue may consist of a large fraction of extra-neurite components.

**Figure 2:**
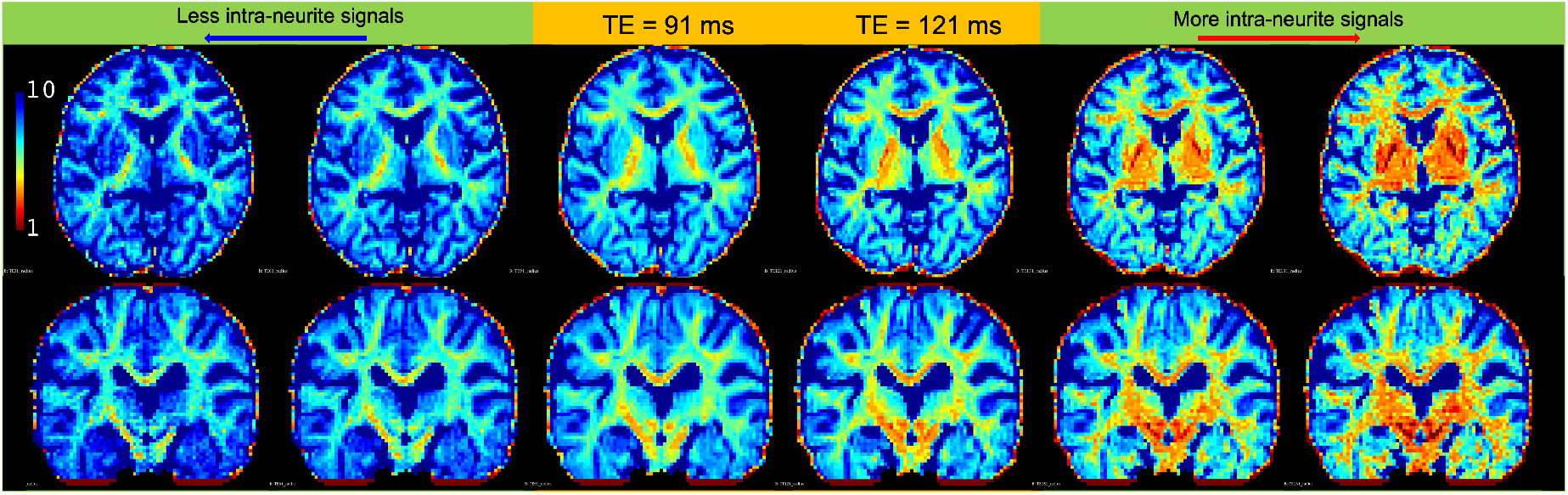
Illustration the estimated effective neurite radius in *μm* at different TEs. The two columns labeled by yellow color show the estimated radius of the acquired data at TE = 91ms, 121 ms, respectively. The two columns on the left show the results with TE = 31ms, 61 ms with reduced volume fraction of intra-neurite signals. The two columns on the right show the results with TE = 151 ms, 181 ms with more signals from intra-neurite space.

Fig. 3 shows the estimated effective neurite radius at TE=211 ms with *r*(*b*) obtained using a constrained NLS method (left panel) and the model in Eq. (17) (right panel). There are several outlier voxels on the left panel which may be related to noisy *r*(*b*) values from the NLS solutions. These outliers are eliminated in the right panel by using the *r*(*b*) function provided by Eq. (17). Most other voxels that are not outliers have similar values between the left and right panels indicating that the model Eq. (17) didn’t induce any systematic biases in the estimation results.

**Figure 3:**
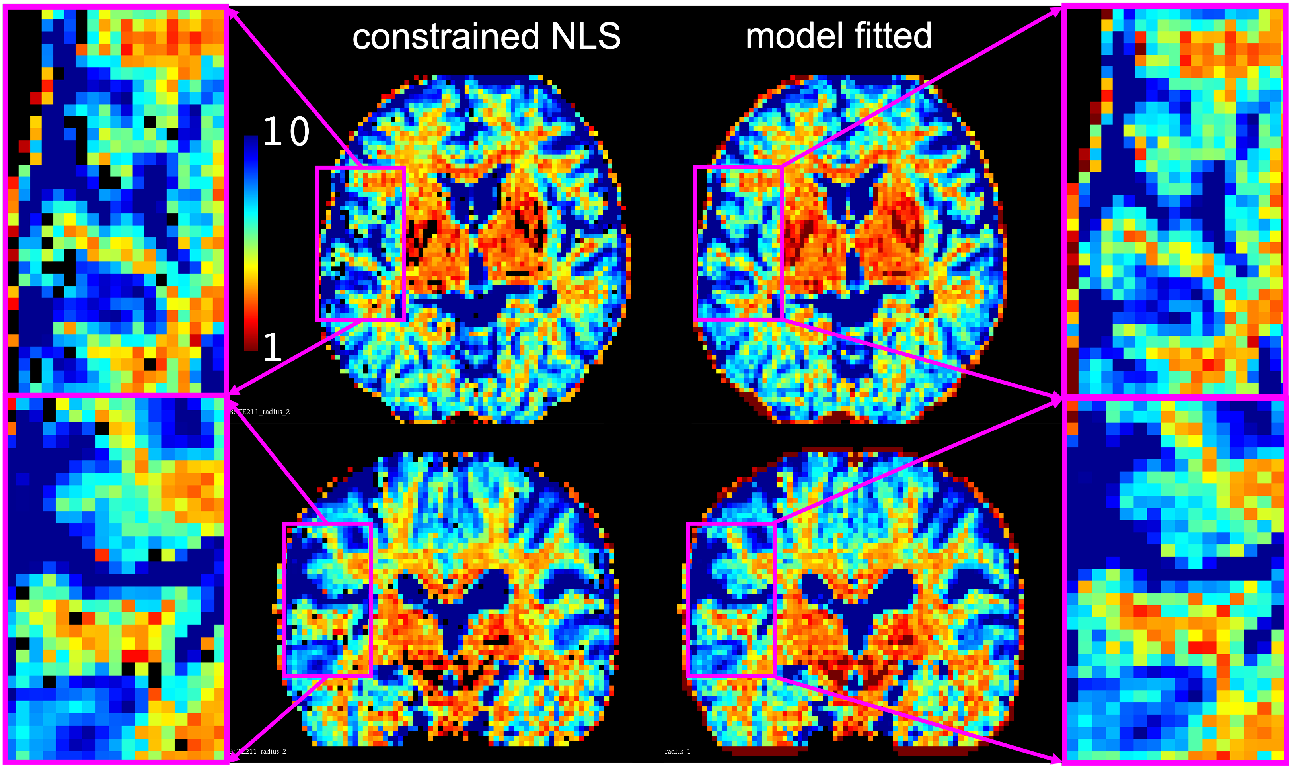
Illustration the estimated effective neurite radius in *μm* at TE = 211 ms. The left panel illustrates the results based on monotonic *r*(*b*) values estimated using a constrained NLS method. The right panel illustrates the results based the model in Eq. (17).

We note that the signal model in Eq. (18) does not restrict *d*_∥_ be ≫ *d*_⊥_. Thereby it may also characterize signals from the cell bodys in gray matter that are not model by sticks. As a result, the estimate neurite size in gray matter also reflects the underlying size of cell body. Results from Figs. 2 and 3 show that the estimated neurite size in the gray matter range from 10 *μm* at TE = 31 ms to 7*μm* at TE = 211 ms which are in good agreement with the size of neural soma of human brains range from 11 ± 7*μm*.

Fig. 4 illustrate the histogram of the estimated neurite radius for all white-matter voxels with fractional anisotropy (FA) > 0.5 with TE=31 ms to TE = 211 ms. The histograms have two modes around 3.8 *μm* and 6 *μm*, respectively. As TE increase, there are less voxels with radii round 6 *μm* and more voxels with radii around 3.8 *μm*.

**Figure 4:**
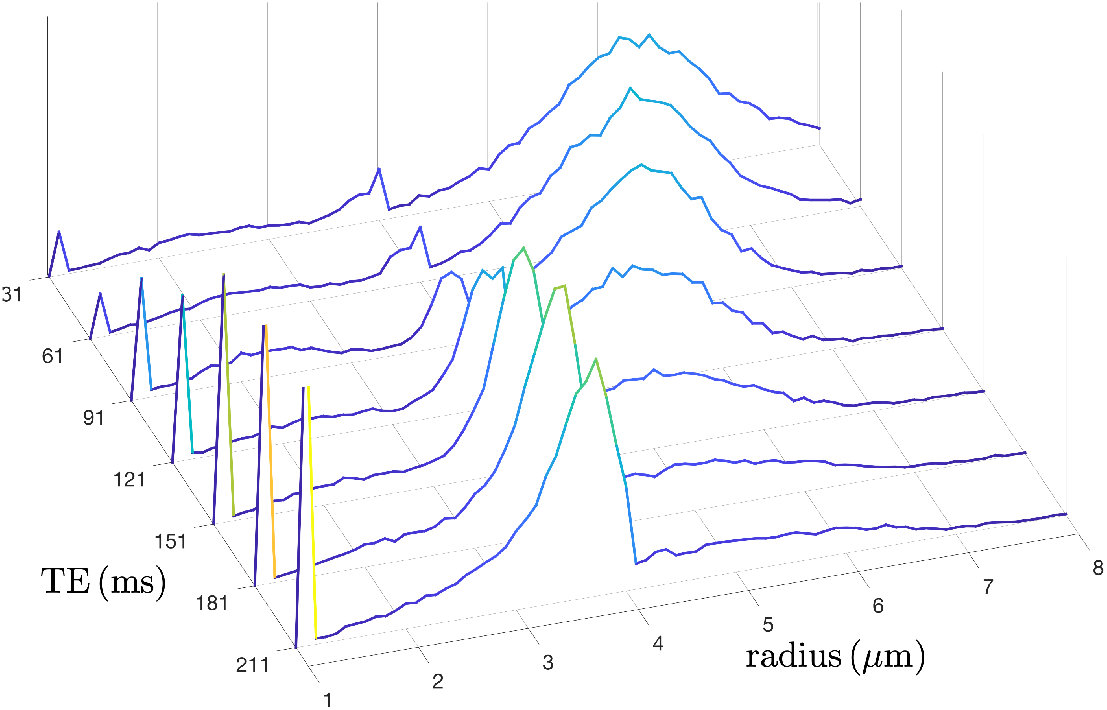
Illustration of the histogram of the neurite radius at different TEs.

## 5 Discussion and conclusion

We have introduced a method for joint modeling of multi-TE diffusion MRI signals for neurite microstructure analysis. Our method exploits the b-value dependent relaxation rates to extrapolate signals to high or low TEs that cannot be directly probed to selectively enhance and attenuate signals from intra-neurite spaces. Thus the the proposed method provide a solution to enhance neurite microstructure estimation without explicitly modeling the extra-neurite signals and fitting multi-compartment signal models. The essence of the proposed method is closely related to the work by [25] which explored ultra-high b-values to suppress the extra-neurite signals. But our method extrapolate signals to high TE values to eliminate extra-neurite signals. Moreover, our method is also closely related to the REDIM techniques proposed in our previous work [17] which uses filters to selectively enhance signals with different relaxation rate and diffusivity to obtain more specific tissue microstructure. Similar to the filters used in [17], the proposed b-value dependent relaxation rate function provides an approach to selectively enhance signals from different components.

Next, we note some limitations and future work. First, acquiring multiple dMRI with multiple TEs increases the scan time. Acquiring more TEs can provide more reliable estimation of the b-value dependent relaxation rates without using a parametric model to reduce noise. But more TEs will prolong the scan time. The rapid time-division multiplexing sequence [9, 10] provides a method to acquire multi-TE dMRI data without significant increasing in scan time. Moreover, advanced denoising methods proposed in [20] may also enhance the reliability of estimation results with fewer samples. Second, the maximum b-value limits the smallest neurite size that can be directly probed. It was shown in [25] that effective neurite radii estimated using b-values upto 25 *ms/μm*^2^ of the white matter are ¡4 *μm* which are smaller and more closer to results from histological studies [1, 6, 14] compared to the proposed results with a mode 4 *μm* at at long TEs. We expect that using higher b-values with a more advanced gradient systems can further improve the estimated neurite size. Thus, in our future work, we will examine the performance of the proposed method with a rapid sequence and advanced MRI system.

In summary, results from this work show that the proposed method can provide more accurate estimation of neurite size and delineate heterogeneous tissue microstructure that cannot be probed by separate analysis of dMRI data with a single TE. Thus, we expect that multi-TE dMRI and the proposed method can provide useful information to investigate tissue abnormality in brain diseases such as epilepsy and other mental disorders.

## Notes

### Competing Interest Statement

The authors have declared no competing interest.

